# Aberrant Information Transfer Interferes with Functional Axon Regeneration

**DOI:** 10.1101/347427

**Authors:** Chen Ding, Marc Hammarlund

## Abstract

Functional axon regeneration requires regenerating neurons to restore appropriate synaptic connectivity and circuit function. To model this process, we developed a single-neuron assay in *C. elegans* that links axon regeneration and synapse reformation with recovery of relevant behavior. After axon injury to the DA9 neuron, regeneration restores synapses to their pre-injury location. Surprisingly, presynapses also accumulate in the dendrite. Both axonal and dendritic synapses are functional. Dendritic synapses result in information misrouting that suppresses behavioral recovery. Formation of dendritic synapses is specifically dependent on dynein-mediated transport and *jnk-1*. However, even when information transfer is corrected, axonal synapses fail to adequately transmit information. Our study reveals unexpected plasticity during functional regeneration. Regeneration of the axon is not sufficient for the reformation of correct neuronal circuits after injury. Rather, synapse reformation and function are also key variables, and manipulation of circuit reformation improves behavioral recovery.

## Introduction

Axon injury disconnects neurons from their post-synaptic targets and destroys circuit functions and behavioral outputs. Axon regeneration—in which injured neurons initiate growth, find appropriate targets, and reconnect functional circuits—has the potential to restore function after nerve injury. Many studies have demonstrated that some behavioral recovery can occur after axon injury. For example, *Caenorhabditis elegans* are able partially to recover locomotion after transection of multiple GABA motor neurons (Byrne et al., 2016; Yanik et al., 2004). Fish recover swimming behavior rapidly following spinal cord injury (Becker et al., 1997; Bernstein, 1964; Briona and Dorsky, 2014; Davis and McClellan, 1993; Oliphint et al., 2010). Further, in many cases, improved recovery after axon injury has been correlated with increased axon regeneration. Increased regeneration in *C. elegans* after treatment with a PARP inhibitor correlates with improved behavioral recovery (Byrne et al., 2016). In the mouse spinal cord, various approaches to improve axon regeneration also result in increased behavioral recovery (Bradbury et al., 2002; Kim et al., 2004; Ramer et al., 2000). Similarly, in the optic nerve, regeneration is increased and partial visual function restored by co-deletion of PTEN and SOCS3 together with application of 4-AP (Bei et al., 2016; Thanos et al., 1997). These studies and others establish a link between axon regeneration and behavioral recovery.

Injured neurons that contribute to behavioral recovery must do so by rewiring into relevant circuits. Consistent with this, regenerated axons have been observed to form new synapses. For example, when injured rat retinal ganglion axons are given permissive conduits that allow regeneration, these neurons can re-establish presynaptic specializations after regenerating into the superior colliculus (Vidal-Sanz et al., 1991; Vidal-Sanz et al., 1987). Similarly, neurons in the optic nerve of goldfish and giant reticulospinal axons of larval lamprey are also able to regenerate synapses (Meyer and Kageyama, 1999; Oliphint et al., 2010; Wood and Cohen, 1981). In the rat peripheral nervous system, regenerated preganglionic axons have also been shown to form new synapses at the originally denervated postsynaptic sites (Raisman, 1977). However, although these morphological and behavioral data suggest that regenerated neurons do form synapses and rewire into circuits, it has not been possible to analyze whether and how individual regenerated neurons contribute to circuit function and behavioral recovery.

Here, we describe a new *in vivo* model for functional regeneration that allows analysis of how individual regenerated neurons rewire into circuits and drive behavior. Using the DA9 motor neuron of *C. elegans,* we show that even after axons regenerate and reform synapses, regenerated neurons recover only a fraction of their original circuit function. A key limitation on functional recovery is misrouted information transfer from regenerated neurons, due to ectopic synapse formation in the dendrite. Preventing ectopic synapse formation by eliminating the MAP kinase *jnk-1* improves functional recovery. Together, our experiments establish the landscape for functional recovery of neurons after regeneration, and show that manipulations that correct the function of regenerated circuits result in improved behavioral recovery after nerve injury.

## Results

### Formation of normal and dendritic synapses after regeneration of a single neuron

DA9 is a bipolar excitatory motor neuron, the most posterior neuron of the DA class, with its cell body in the ventral midline. According to the electron microscopy reconstruction, the DA9 dendrite extends anteriorly in the ventral nerve cord (VNC), receiving input from command neurons and sensory neurons (Hall and Russell, 1991; White et al., 1976; White et al., 1986) (Figure 1A). The DA9 axon extends posteriorly in the VNC and then crosses to the dorsal nerve cord (DNC) via a commissure. It then proceeds anteriorly in the DNC and forms en passant cholinergic synapses that are dyadic with the posterior dorsal body wall muscles (BMWs) and the ventral D (VD) inhibitory GABA motor neurons. These synapses are restricted to a small region of the DA9 axon, and the axon itself extends in the DNC past the synaptic region. Based on its anatomy and connectivity, DA9 activity is predicted to trigger dorsal bending of the animal’s tail region, by directly activating dorsal BMWs while simultaneously inhibiting opposing ventral BMWs via VD activity.

**Figure 1.**
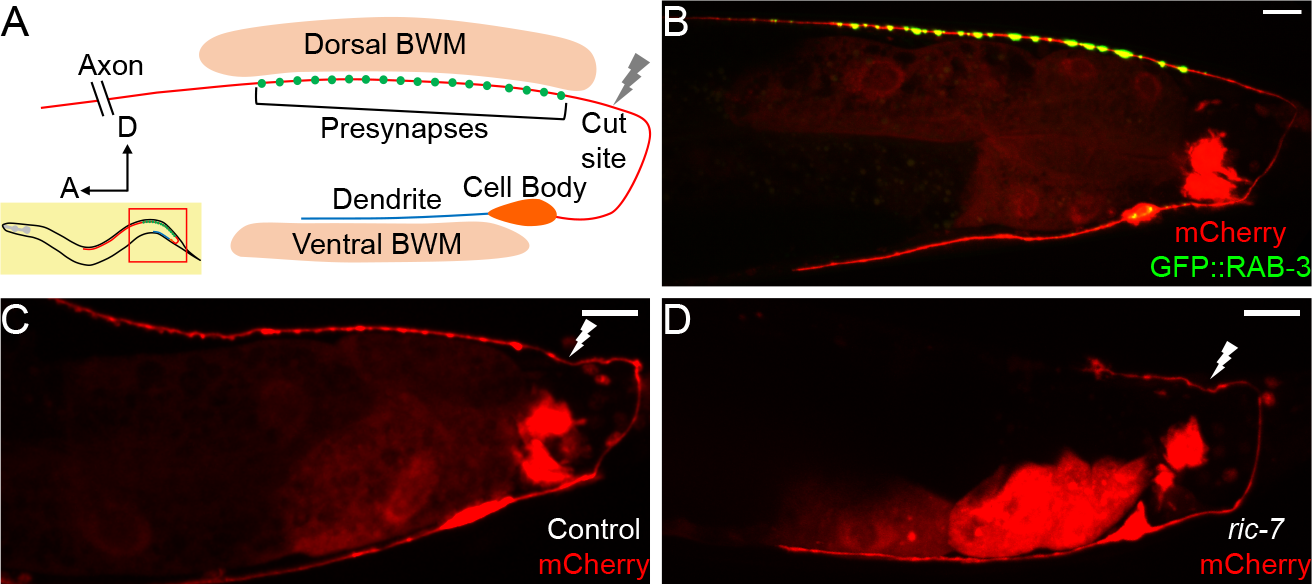
DA9 axon regeneration and degeneration. (A) Diagram of DA9 morphology and the laser cutting site. (B) A control adult expressing GFP::RAB-3 and mCherry in DA9. mCherry expression is also observed in the posterior gut in these micrographs. (C) A control animal 12hr post-axotomy. The distal axon fragment remains and overlaps with the regenerating axon. The white lightning mark indicates the axotomy site. (D) A *ric-7* animal 12hr post-axotomy. The distal fragment has degenerated. Scale bars = 10 μm.

To analyze the morphology of DA9 neurites and synapses in the context of axon regeneration, we used the DA9-specific *itr-1 pB* promoter to co-express soluble mCherry along with the presynaptic marker GFP-RAB-3, as previously described. In the posterior of the animal, this promoter drives expression exclusively in the DA9 neuron, as well as in the posterior cells of the intestine (Klassen and Shen, 2007) (Figure 1B). These labels allowed us to observe the DA9 dendrite, axon, and synaptic vesicle clusters, and were consistent with previous observations and with the electron microscopy reconstruction (Hall and Russell, 1991; Klassen and Shen, 2007; White et al., 1976). We used a pulsed laser essentially as described (Byrne et al., 2011) to sever the DA9 axon just posterior to its synaptic region in the DNC. We severed axons in L4 stage animals and assessed regeneration. We found that 12 hours after axotomy, DA9 axons had initiated regeneration and regenerated past the injury site (Figure 1C), similar to the kinetics of regeneration in other neurons (Chuang et al., 2014; Hammarlund and Jin, 2014; Yanik et al., 2004). However, even after 12 hours, the distal axon fragment still was present (Figure 1C), similar to slow removal of fragments after injury in other *C. elegans* neurons (Nichols et al., 2016). Presence of this distal fragment raised the possibility that regenerating DA9 axons might restore connectivity simply by fusion with the fragment, as previously observed in *C. elegans* (Abay et al., 2017; Ghosh-Roy et al., 2010; Neumann et al., 2015; Neumann et al., 2011). Alternatively, the fragment might interfere with synaptic regeneration by occupying relevant anatomical sites. By contrast, distal axon fragments degenerate rapidly in vertebrates and *Drosophila* through a process called Wallerian degeneration (MacDonald et al., 2006; Martin et al., 2010; Waller, 1850). In these cases, successful degeneration is thought to be permissive for axons to regenerate and re-innervate the targets. Indeed, delaying Wallerian degeneration has been shown to impair regeneration and delay the locomotor recovery of mice (Bisby and Chen, 1990; Brown et al., 1994; Zhang et al., 1998).

In order to build a permissive environment for synaptic regeneration and block fusion, we used a recently identified mutation, *ric-7* (*n2657*), to accelerate degeneration of DA9 distal axon fragments after injury (Hao et al., 2012; Nichols et al., 2016; Rawson et al., 2014). We found that DA9 developed normally in *ric-7* animals compared to controls (Figure 1 supplement 1A-1D), consistent with previous data (Hao et al., 2012). However, after laser axotomy of DA9, the distal axon segment degenerated quickly and was largely cleared away at 12 hours (Figure 1D). In the GABA motor neuron system, *ric-7* animals also show enhanced degeneration together with increased regeneration 24 hours after axotomy, indicating that loss of *ric-7* and enhanced degeneration does not interfere with axon regeneration (Figure 1 supplement 1E-1H). These experiments indicate that *ric-7*(*n2657*) eliminates potential interference from the stubborn distal axon fragments, facilitating the study of new synapse formation during axon regeneration. Therefore, we included the *ric-7* mutation in the following experiments unless further mentioned.

We characterized the characteristics and kinetics of DA9 axon regeneration and synapse reformation by examining animals at different time points after axotomy (Figure 2A-2D). We found that axon growth occurs only in the first 48 hours, after which no additional growth occurs (Figure 2G). During this time, regenerating axons completely reinnervate their previous synaptic region, but fail to grow far past the synaptic region as observed before injury (Figure 2D, 2G; the grey stripe in Figure 2G indicates the synaptic region of regenerated axons 48hr post-axotomy). Lack of growth into the distal asynaptic area results in regenerated axons that only reach 1/3 of length of intact axons (Figure 2G, cut 72hr vs uncut 48hr, mean=182.0 μm and 518.2 μm, p<0.0001, unpaired t test). Overall, 100 percent (28/28) of severed DA9 axons initiated regeneration and reinnervated the synaptic region. The high incidence of regeneration and accurate growth to the former synaptic area we observed is ideal for analyzing the function of regenerated neurons, and is likely due to our choice of injury site close to the DNC. By contrast, if DA9 is severed at the dorsal-ventral midline, it often fails to reinnervate the synaptic area (data not shown).

**Figure 2.**
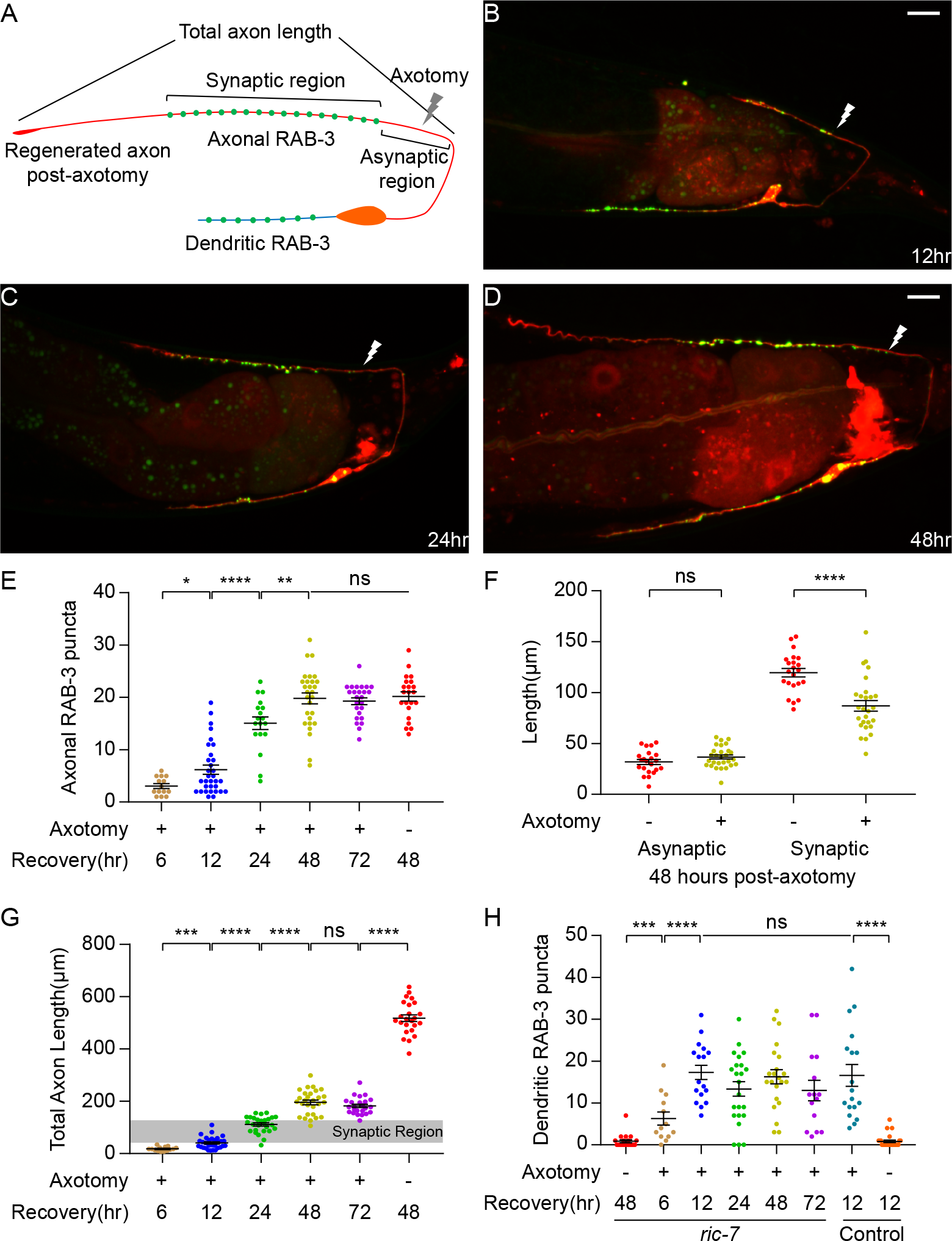
DA9 forms normal and dendritic synapses during regeneration. (A) Diagram of a regenerated DA9 axon. (B-D) DA9 axonal and synaptic regeneration at different timepoints post-axotomy. The white lightning mark indicates the axotomy site. Scale bars = 10μm. (E) Number of axonal RAB-3 puncta at different timepoints with or without axotomy. Mean + SEM. *p<0.05; **p<0.01; ****p<0.0001; ns, not significant. Unpaired t test for comparisons between two groups and one-way ANOVA for more than two groups. (F) Length of the asynaptic and the synaptic region in intact and axotomized animals 48hr post-axotomy. Mean + SEM. ****p<0.0001; ns, not significant. Unpaired t test. (G and H) Total axon length and number of dendritic RAB-3 puncta at different timepoints with or without axotomy. Mean + SEM. ***p<0.001; ****p<0.0001; ns, not significant. Unpaired t test for comparisons between two groups and one-way ANOVA for more than two groups.

We found that as DA9 regenerates, it forms new SV puncta coincident with regeneration (Figure 2B-2E). A prominent feature of DA9 synapses in intact neurons are their stereotypic placement along the DNC, in which there is a specific synaptic region containing a stereotyped number of puncta that occupies a limited and stereotyped region of the axon (Figure 1A, 1B) (Hall and Russell, 1991; Klassen and Shen, 2007; Poon et al., 2008). Regenerated neurons were largely able to restore the stereotyped arrangement of SV puncta. By 48 hours, when axon growth is completed (Figure 2G), the number of SV puncta was statistically indistinguishable from intact animals and did not further increase (Figure 2E, cut 48hr, cut 72hr and uncut 48hr, mean=19.8, 19.3 and 20.2, p=0.7864, one-way ANOVA). The new synapses are again localized in a specific region of the DNC (Figure 2D). The position of the new synaptic region, as indicated by the length of the asynaptic region (Figure 2A), is similar to intact controls (Figure 2F, uncut vs cut, mean=31.8 μm and 36.7 μm, p=0.1238, unpaired t test). The large difference is in the length of the synaptic region, which is shorter than in uninjured axons (Figure 2F, uncut vs cut, mean=119.5 μm and 87.0 μm, p<0.0001, unpaired t test). Overall, these data indicate that the regenerating DA9 axon is able to form new SV puncta and largely re-establish its synaptic pattern. The process of axon regeneration, both in terms of axon regrowth and synapse reformation, has ended by 48 hours after axotomy.

Strikingly, we found that SV puncta also form in the dendrite during regeneration (Figure 2A-2D). In intact animals, SV puncta were essentially absent from the dendrite (Figure 1B). After axon surgery, however, dendritic SVs became apparent at as early as 6 hours after axotomy (Figure 2H). The number of puncta reached a maximum at 12 hours (mean=17.3) and decreased only slightly thereafter. Importantly, the appearance of ectopic SVs in the dendrite is not caused by the *ric-7* background, because control animals also accumulate a similar number of ectopic puncta in response to axotomy (Figure 2H, *ric-7* cut 12hr vs control cut 12hr, mean=17.3 and 16.6). These data show that axonal injury triggers ectopic accumulation of SVs in the dendrite, identifying an unexpected form of plasticity during recovery from nerve injury.

### Aberrant functional recovery of a single neuron-driven behavior

To determine the effects on function of these morphological changes in DA9 after axotomy, we sought to establish a behavioral assay entirely dependent on DA9 synaptic output. We expressed Chrimson, a red-shifted channelrhodopsin, in DA9 using the DA9-specific *itr-1 pB* promoter and activated it with green light (Klapoetke et al., 2014; Schild and Glauser, 2015). The anatomy of DA9 predicts that it will activate the posterior dorsal BMWs and inhibit the posterior ventral BMWs, causing the tail to bend dorsally (Figure 3A). Indeed, when 5 seconds of light stimulation was applied, freely-behaving Chrimson-expressing animals displayed strong dorsal tail bending (Figure 3B). We quantified this behavior by measuring the tail angle using the WormLab software package. Animals that expressed Chrimson and were cultured with all-trans-retinal (ATR) showed an average of 80 degrees of dorsal bending, time-locked to the light stimulus (Figure 3C). Dorsal bending in response to light was observed in every animal (N=42). After the stimulus ended, the tail gradually returned to its normal position. Animals that did not express Chrimson, or that were not given ATR, showed no response to light stimulation (Figure 3 supplement 1A, 1B and 1D). In addition, the behavior of Chrimson-expressing animals cultured with ATR was abolished 12 hours after DA9 cell body was ablated by laser (Figure 3 supplement 1C and 1D). Thus, this assay allows the precise analysis of the ability of a single neuron to drive behavior, and (in intact animals) has very low levels of inter-animal variability.

**Figure 3.**
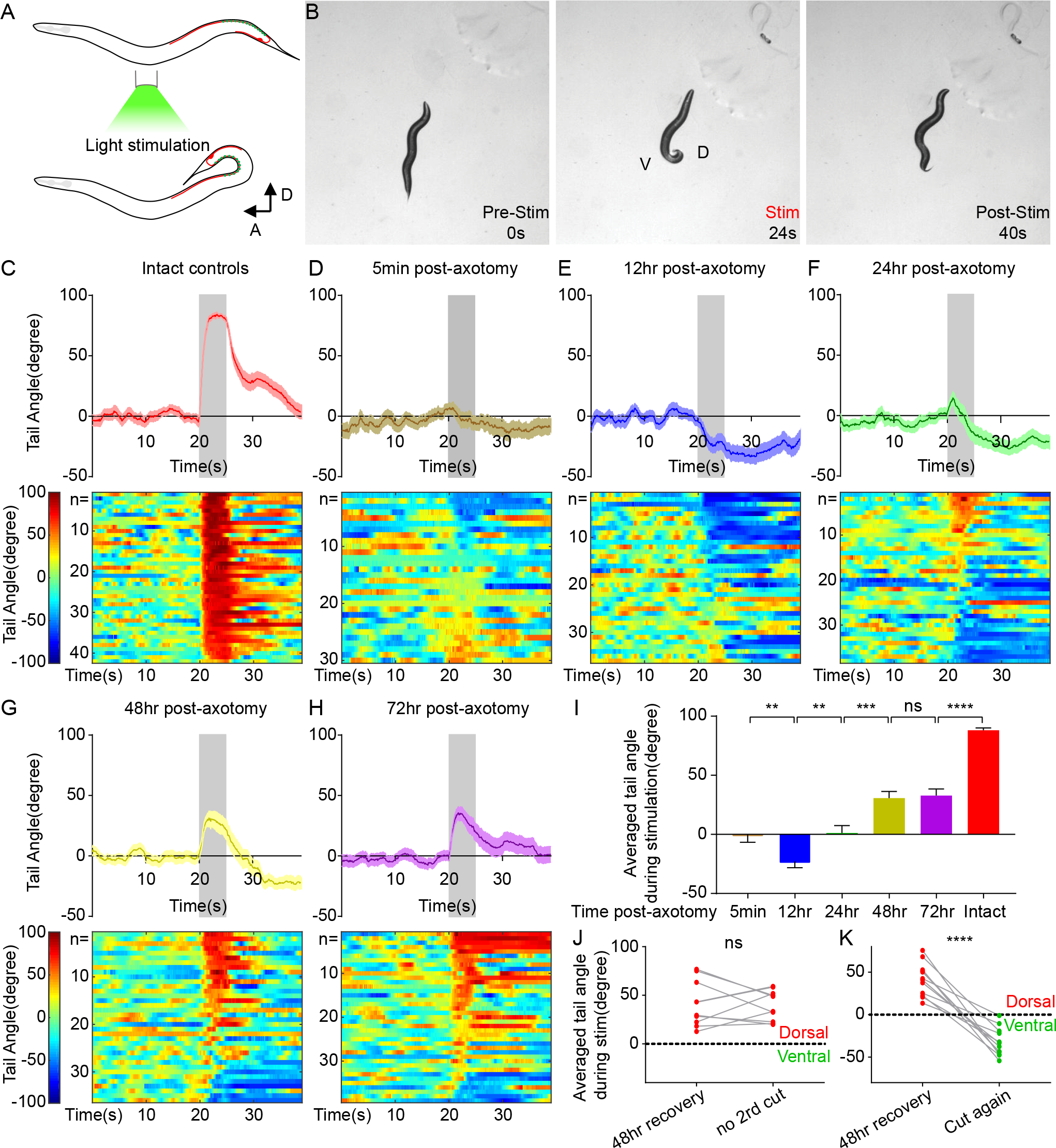
Aberrant functional recovery of a single neuron-driven behavior. (A) Diagram of the optogenetically triggered tail-bending behavior. (B) Images from an example movie showing the dorsal tail-bending behavior. (C-H) Quantification of the tail-bending behavior of intact controls (C), axotomized animals 5min post-axotomy (D), 12hr post-axotomy (E), 24hr post-axotomy (F), 48hr post-axotomy (G) and 72hr post-axotomy (H). Both averaged data (upper, Mean + SEM) and heat maps (lower) of individual responses are shown. Shaded areas represent the 5 second light stimulation. The color bar indicates the tail angle. Positive numbers indicate dorsal bending, and negative numbers indicate ventral bending. (I) Averaged tail angle during the 5 second stimulation at different timepoints post-axotomy of animals in (C-H). Mean and SEM. **p<0.01; ***p<0.001; ****p<0.0001; ns, not significant. Unpaired t test. (J and K) Recovery of dorsal bending is abolished 12hr after the second cut of the regenerating axon (K) but stays the same in the absence of the second cut (J). ****p<0.0001; ns, not significant. Paired t test.

We then examined the ability of DA9 to mediate tail bending after acute axon injury. 5 minutes after axotomy of DA9, light activation of DA9 did not cause dorsal bending behavior (Figure 3D). These data confirm that the bending behavior is specifically driven by DA9, and indicate that axotomy proximal to the known DA9 synaptic region completely disconnects DA9 from its outputs. Even though the distal fragment containing the synaptic puncta contained Chrimson and was exposed to light (together with the rest of the animal), this was insufficient to trigger tail bending. Thus, Chrimson-mediated neuronal output in DA9 requires connection to the cell body.

48 hours after axon injury, when axon and synapse regeneration is complete (Figure 2E, 2G), we found that behavior has partially recovered (Figure 3G). At 72 hours, consistent with the lack of further morphological growth after 48 hours, there was no further behavioral recovery (Figure 3H). Thus, the maximum behavioral recovery is on average about 1/3 of the control animals (Figure 3I, cut 72hr vs control, mean=26.3° and 70.5°, p<0.0001, unpaired t test). A major reason for limited recovery on average was that recovered behavior after injury is far more variable than behavior in intact animals: while more than half of injured animals recovered dorsal bending, many did not. The variability in functional recovery is in sharp contrast to the robust morphological axon regeneration and synapse reformation we observed (Figure 2E, 2G), suggesting that rebuilding a functional circuit requires additional biological factors beyond what can be assessed using morphological analysis of the axon. Nevertheless, most animals did recover behavior, indicating that newly formed SV puncta in the axon are functional regenerated synapses. To confirm that the behavior recovery is indeed a result of synaptic regeneration of DA9 (rather than some other form of plasticity), we recut DA9 in animals that displayed behavioral recovery at 48 hours, and measured the behavior 12 hours later, before a second round of regeneration could occur. Recovered dorsal bending was completely abolished after recutting, indicating that behavioral recovery is due to DA9 regeneration (Figure 3J, 3K). Together, these experiments indicate that the single-neuron behavioral assay reveals the functional output of DA9 after axon regeneration.

We also analyzed behavior at earlier time points, when SV puncta are at their maximum in the dendrite and axonal regeneration is not yet complete (Figure 2B). At 12 hours after axotomy, we found that on average, DA9 activation results in ventral rather than dorsal bending (Figure 3E, 3I). Data from individual animals reveal that at 12 hours, more than half of the animals bend their tail ventrally, rather than dorsally. Thus, DA9 activation in these regenerated neurons is causing behavior opposite to uninjured neurons. This aberrant behavior is observed less frequently as regeneration proceeds, but even at 48 and 72 hours after injury some animals still exhibit ventral bending (Figure 3G, 3H). These data demonstrate that although injury and regeneration can restore normal behavior in some individuals, in other individuals injury and regeneration result in formation of a pathological circuit that drives novel and inappropriate behavior.

### Rerouted information transfer in a regenerated circuit

Aberrant ventral bending behavior is maximal at the same time that SV puncta in the dendrite are maximal. This observation suggests a cell-biological explanation for aberrant behavior after regeneration: that the ectopic dendritic SVs are capable of releasing neurotransmitters on the ventral side and activating the ventral BWMs, which normally receive input from other neurons in the ventral nerve cord (Figure 1A). As more new SV puncta are added to the dorsal axon later, release at the dorsal side eventually predominates (Figure 3E-3H). To test this hypothesis and to better understand functional regeneration and circuit plasticity, we combined optogenetic stimulation and Ca^2+^ imaging to monitor the activity of postsynaptic BWMs during DA9 regeneration. This approach allowed us to monitor Ca^2+^ levels in both dorsal and ventral BWMs simultaneously during activation of DA9. For this analysis, we focused on Ca^2+^ level changes during DA9 stimulation.

In uncut controls, we found that dorsal BWMs displayed robust increases in Ca^2+^ levels during light stimulation of DA9, consistent with DA9 directly activating these muscles (Figure 4A, 4C). We also observed simultaneous decreases in ventral BWM Ca^2+^ levels, presumably mediated by DA9 activation of inhibitory GABAergic VD motor neurons as predicted by the wiring diagram (Figure 4F, 4J). These effects on the muscles should trigger dorsal muscle contraction and ventral relaxation, resulting in the dorsal bending we observed in our behavioral analysis. Thus, simultaneous stimulation and calcium imaging allows interrogation of functional connectivity in this simple motor circuit.

**Figure 4.**
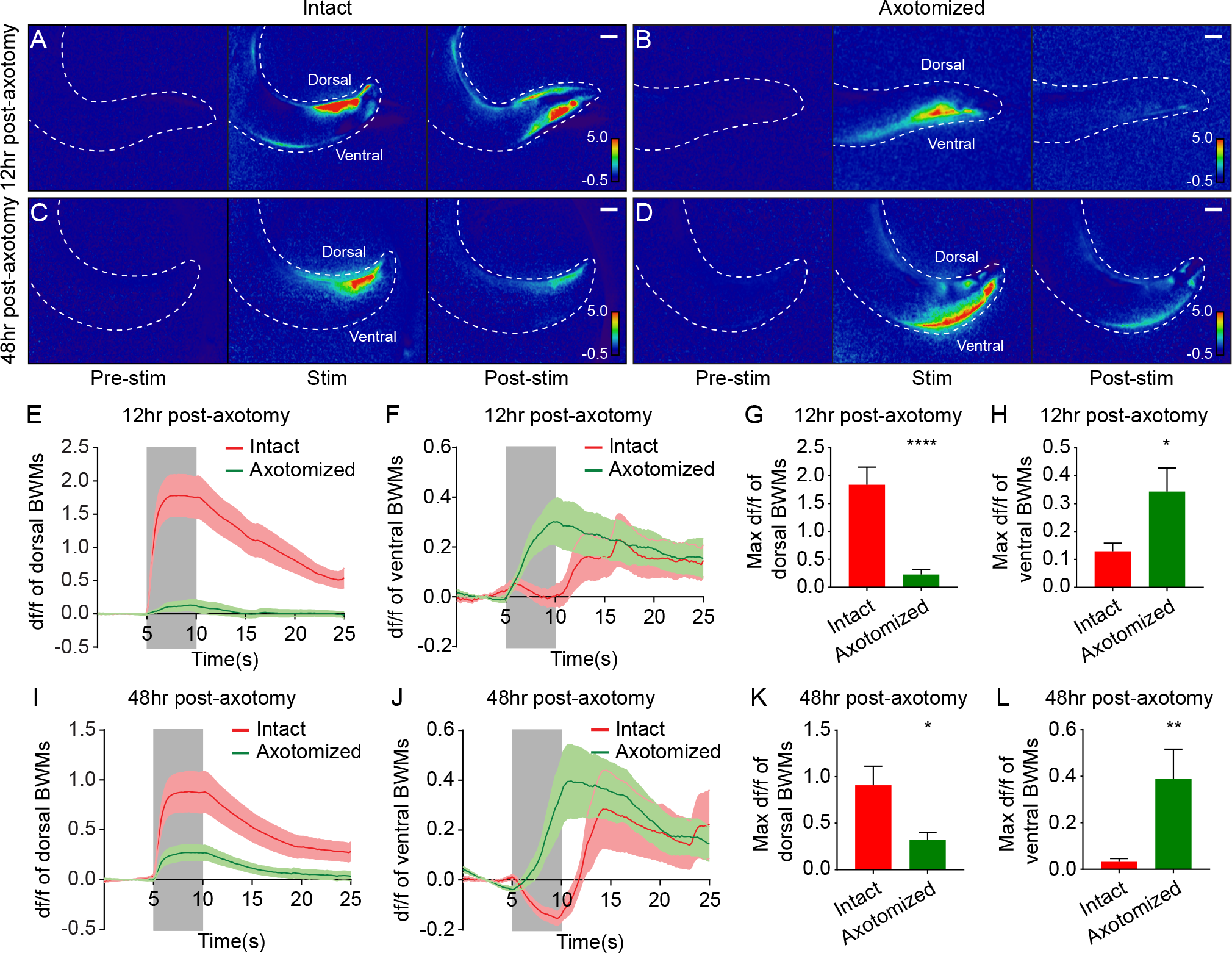
Simultaneous optogenetic stimulation and calcium imaging reveals rerouted information transfer in the regenerated circuit. (A-D) Heat maps of calcium signal fold changes (df/f) in controls (A and C) and axotomized animals (B and D) 12hr or 48hr post-axotomy. White dashed lines delineate the animals’ tails. Scale bars = 20 <m. (E-H) Calcium signal traces (E and F) and peak amplitudes during stimulation (G and H) of both dorsal and ventral BWMs in controls and axotomized animals 12hr post-axotomy (n= 29 for axotomized animals and 25 for intact controls). Shaded areas indicate the 5 second light stimulation. Mean + SEM. *p<0.05; ****p<0.0001. Unpaired t test. (I-L) Calcium signal traces (I and J) and peak amplitudes during stimulation (K and L) of both dorsal and ventral BWMs in controls and axotomized animals 48hr post-axotomy (n= 30 for axotomized animals and 31 for intact controls). Shaded areas indicate the 5 second light stimulation. Mean + SEM. *p<0.05; **p<0.01. Unpaired t test.

In sharp contrast to the results in uninjured animals, we found that 12 hours after axotomy, dorsal BWMs showed little Ca^2+^ increase in response to light stimulation (Figure 4B, 4E). The maximum Ca^2+^ amplitude in dorsal BWMs 12 hours after axotomy was about 12% of the control animals (Figure 4G, mean=0.23 and 1.84 with and without axotomy). Further, DA9 activation 12 hours after axotomy drove strong Ca^2+^ increases in ventral BWMs, consistent with the dendritic SV localization and the ventral bending behavior at the 12 hour time point (Figure 4F, 4H). 48 hours after axotomy, DA9 activation was more effective at driving dorsal BWMs, which displayed Ca^2+^ increases about 35% of the controls, confirming that new synapses in the axon are functional (Figure 4D, 4J, 4K; mean=0.32 and 0.91 with and without axotomy in Figure 4K). Even at the 48 hour time point, however, the ventral BWMs also displayed Ca^2+^ increases, suggesting that the ectopic synapses in the dendrite are still functional at 48 hours after axotomy (Figure 4J, 4L). Together, these data demonstrate that information transfer—the fundamental role of neurons—is rerouted after axon injury and regeneration.

### Aberrant information transfer is independent of *dlk-1* and suppresses behavioral recovery

Even when regeneration is complete and axonal synapses have reformed (at 48 and 72 hours), recovery of the ability of DA9 to drive behavior is only partial. We hypothesized that at least part of this deficit is due to aberrant information transfer at these later time points. Specifically, release of SVs from the ectopic synapses in the dendrite would be expected to suppress dorsal bending of the tail, by exciting ventral BWMs and also by exciting DD GABA neurons that would in turn inhibit dorsal BWMs according to the wiring diagram (Schuske et al., 2004). We tested the ability of dendritic release to drive behavior at these late time points by using mutants in *dlk-1*, a MAPKKK that is a key regulator of regeneration in *C. elegans* and other species (Hammarlund et al., 2009; Shin et al., 2012; Xiong et al., 2010; Yan et al., 2009). Consistent with these previous results, we found that DA9 axons completely fail to regenerate in *ric-7; dlk-1* animals (Figure 5A-5C). Despite this lack of axonal response, however, *dlk-1* mutant animals still mislocalize SV puncta ectopically to the dendrite after axotomy, at similar levels as controls (Figure 5B, 5D). Thus, the *dlk-1* background enables analysis of the function of the dendritic SVs in the absence of the SVs in the regenerating axon. Loss of *dlk-1* had no detectable effect on the morphology (Figure 5A) or function of DA9; in the absence of injury, *dlk-1* mutant animals showed robust dorsal bending of the tail (Figure 5 supplement 1A, 1C). However, when DA9 is activated in *dlk-1* mutant animals 12 or 48 hours after axotomy, almost all animals bend their tail ventrally (Figure 5E, Figure 5 supplement 1B-1D). Together these data indicate that ectopic synapses in the DA9 dendrite can mediate ventral bending even at 48 hours after axotomy. These data also indicate that mislocalization of SVs to the dendrite in response to injury is independent of *dlk-1* function.

**Figure 5.**
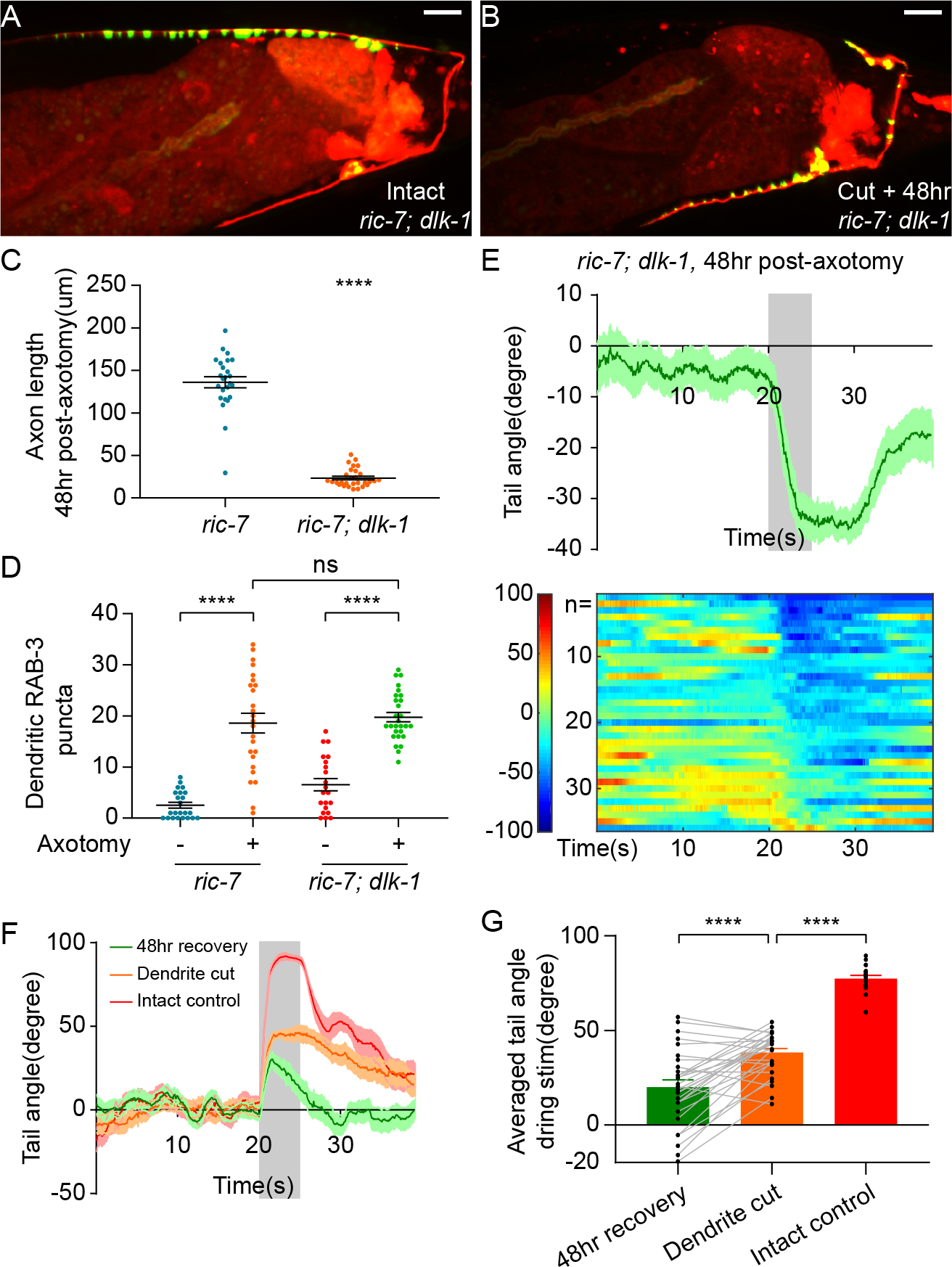
Aberrant information transfer is independent of dlk-1 and suppresses behavioral recovery. (A and B) GFP::RAB-3 and mCherry labeling of DA9 in intact and axotomized *ric-7; dlk-1* animals 48hr post-axotomy. Scale bars = 10μm. (C and D) Comparison of length of regenerating axons (C) and number of dendritic RAB-3 puncta (D) between *ric-7* and *ric-7; dlk-1* animals 48hr post-axotomy. Mean + SEM. ****p<0.0001; ns, not significant. Unpaired t test. (E) Ventral tail-bending behavior of *ric-7; dlk-1* animals 48hr post-axotomy. The shaded area indicates the 5 second light stimulation. Mean + SEM. (F) Traces of tail-bending behavior of *ric-7* intact controls (red) and axotomized animals (green) 48hr post-axotomy. Dendrites of axotomized animals were then cut and behavior was measured again 2 hours later (orange). Mean + SEM. The shaded area indicates the 5 second light stimulation. (G) Averaged tail angles during stimulation of the animals in (F). Mean and SEM. ****p<0.0001. Paired t test between 48hr recovery and dendrite cut. Unpaired t test between dendrite cut and intact controls.

To test the idea that aberrant dendritic SV release suppresses behavioral recovery after axotomy, we eliminated dendritic release by removing the dendrite after axon regeneration was complete. 48 hours after axotomy, we assessed behavior, severed the dendrite, and assessed behavior again 2 hours later. We found that removing the dendrite increased dorsal bending about twofold (Figure 5F, 5G).

However, even after dendrite removal, DA9-driven behavior still did not reach the level observed in uninjured controls (Figure 5G, 48hr recovery vs dendrite cut vs control, mean=20.0, 38.2 and 77.4, paired t test between 48hr recovery and dendrite cut, unpaired t test between dendrite cut and control). Together, these results indicate that even after regeneration is complete, misdirected SV release from the dendrite suppresses behavioral recovery. Further, these data indicate that axonal activity, despite largely normal synaptic morphology (Figure 2E, 2F), is only able to drive behavior to about 50% of pre-injury levels even when dendritic suppression is removed.

### Dendritic microtubule polarity is maintained after axotomy

We next sought to understand the mechanism of ectopic synapse formation in the dendrite. In some cases, axonal injury to a neuron can cause one of its dendrites to become more axon-like (Dotti and Banker, 1987; Gomis-Ruth et al., 2008; Stone et al., 2010). This conversion can be accompanied by a change in microtubule (MT) polarity (Stone et al., 2010; Takahashi et al., 2007). MTs in *Caenorhabditis elegans* axons are uniformly polarized with their plus-ends oriented away from the cell body, while MTs in dendrites are largely oriented with minus-ends facing the distal tip (Maniar et al., 2011; Yan et al., 2013). If axon injury to DA9 results in a change in MT polarity, so that dendritic MTs become axonlike with plus-ends out, this polarity change could account for SV mislocalization to the DA9 dendrite after axotomy. We examined microtubule polarity in DA9 using live imaging of fluorescently tagged MT plus-end-tracking protein EBP-2, the *C. elegans* homolog of human EB1 (Srayko et al., 2005; Yan et al., 2013) (Figure 6A-6E). To examine the effect of injury, imaging was done 12 hours after injury, a time point when dendritic SVs were abundant (Figure 2H). We found that in uninjured DA9 neurons, MT polarity in the axon was nearly completely plus-end out, while MT polarity in the dendrite was dominated by plus-end in MTs, consistent with previous results (Maniar et al., 2011; Yan et al., 2013) (Figure 6F). Surprisingly, however, axon injury did not result in polarity reversal in the axon or dendrite; in fact, in the dendrite after axon injury plus-end in MTs were even more dominant than before injury (Figure 6F).

**Figure 6.**
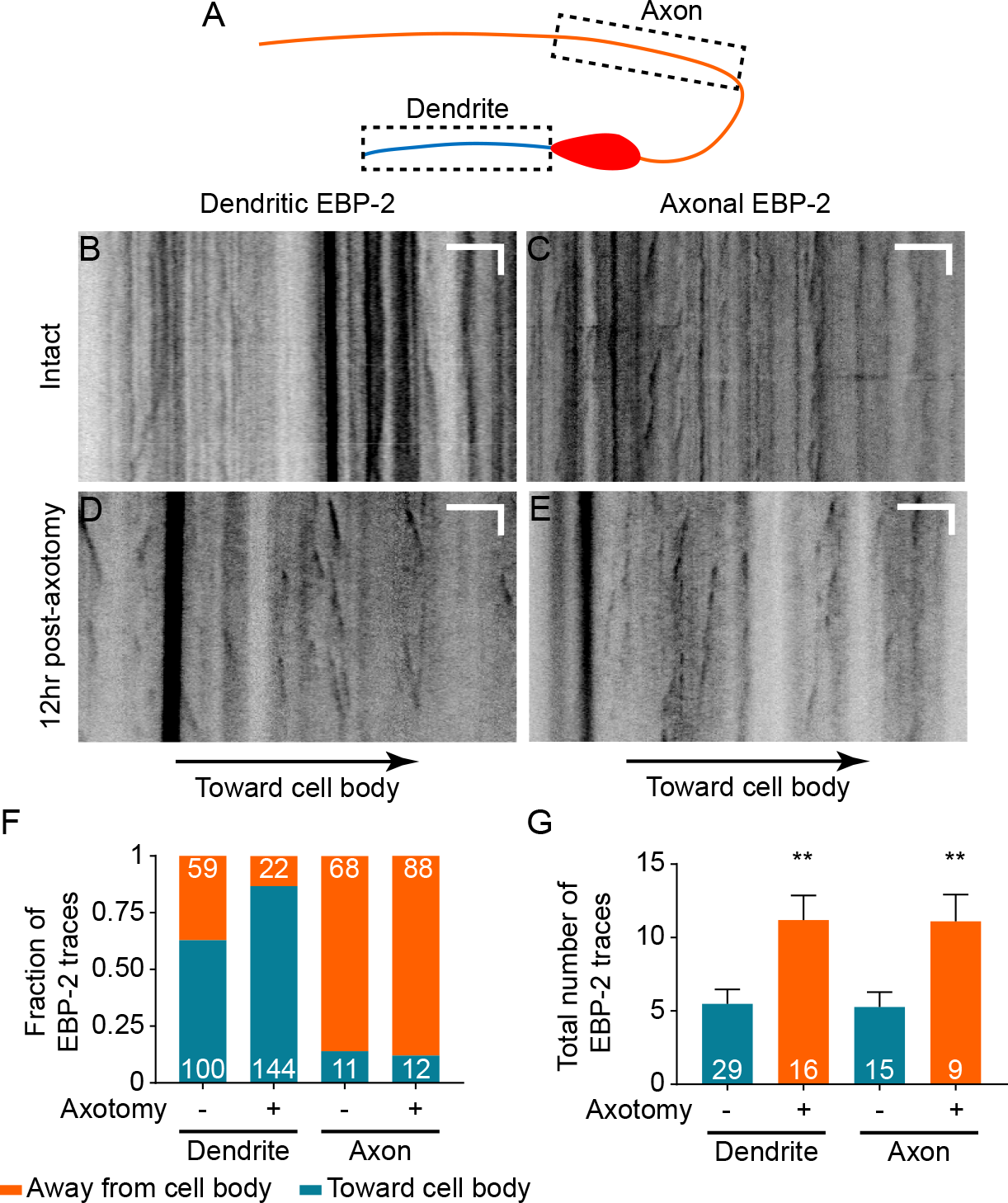
Dendritic microtubule polarity is maintained after axotomy. (A) Diagram of regions of EBP-2::GFP imaging in DA9. (B-E) Kymographs of EBP-2 traces in both DA9 dendrite and axon in intact and axotomized animals 12hr post-axotomy. Scale bars are 5μm and 5s. (F) Direction of EBP-2 traces in DA9 dendrite and axon. Dendritic microtubule polarity is opposite to axonal microtubule polarity. Numbers represent the number of EBP-2 traces. (G) Total number of EBP-2 traces in DA9 dendrite and axon with and without axotomy. Numbers represent the number of animals. Mean and SEM. **p<0.01. Unpaired t test.

Although axotomy did not reverse the polarity of MTs, it did have a significant effect on MT dynamics in both axons and dendrites. In both compartments, we observed an increased number of EBP-2 traces (Figure 6G), which has also been observed in *C. elegans* mechanosensory neurons (Chuang et al., 2014; Ghosh-Roy et al., 2012) and *Drosophila* sensory neurons (Stone et al., 2010). Overall, neuronal injury in DA9 results in increased MT dynamics but no change in MT polarity. Thus, dendritic SV mislocalization is not a consequence of MT polarity changes.

To further test the idea that axon injury might convert the DA9 dendrite into an axon-like process, we examined the localization of the dendritic nicotinic acetylcholine receptor (nAChR) subunit *acr-2* (Squire et al., 1995). We found that in uninjured DA9 neurons, ACR-2::GFP is highly enriched in the dendrite and soma and is absent from the axon (Figure 6 supplement 1A), as previously described (Barbagallo et al., 2010; Qi et al., 2013; Yan et al., 2013). 12 hours after injury, when ectopic SVs are highly enriched in the dendrite (Figure 2H), localization of ACR-2::GFP was not different from uninjured neurons (Figure 6 supplement 1B). Together, these results indicate that axon injury does not globally push the DA9 dendrite toward becoming axon-like, and suggest that more specific cellular mechanisms must mediate ectopic SV localization.

### Dynein-mediated transport is required for SV mislocalization to the dendrite after injury

Previous studies have shown that UNC-104/KIF1A is the primary motor responsible for transporting SVs towards the MT plus-end (Hall and Hedgecock, 1991; Okada et al., 1995; Pack-Chung et al., 2007), while the cytoplasmic dynein complex is involved in MT minus-end directed transport in neurons (Goldstein and Yang, 2000; Koushika et al., 2004). Since MT polarity is maintained after axotomy, so that dendritic MTs are primarily plus-ends in, we reasoned that ectopic synapse formation in the dendrite may be mediated by the minus-end-directed dynein motor complex. To test this, we examined SV puncta in the dendrite in *dhc-1(js319)* mutant animals (Koushika et al., 2004). These mutants had largely normal SV distribution in DA9 prior to injury (Figure 7 supplement 1A, 1B). However, after injury we found that the *dhc-1(js319)* mutation completely suppresses the dendritic accumulation of SVs at 48 hours after axotomy (Figure 7B, 7E). Partial suppression was obtained in animals mutant for another component of the dynein complex, *nud-2*(*ok949*), which is the worm ortholog of Nudel (Figure 7C, 7E). Together these data indicate that dynein mediated transport is required for SV mislocalization in the dendrite in response to axotomy.

**Figure 7.**
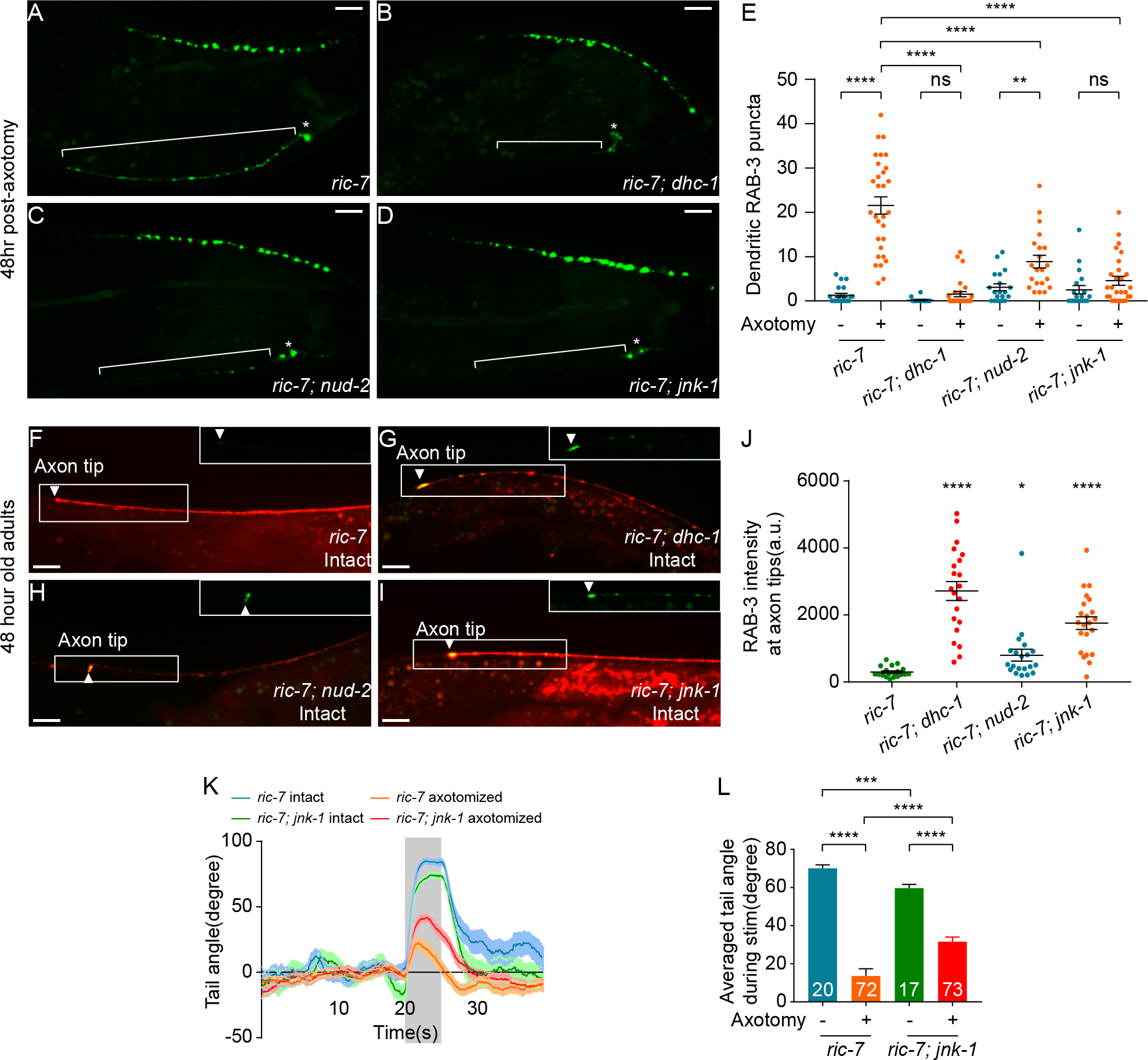
JNK-1 and dynein-mediated transport mediate SV mislocalization to the dendrite. (A-D) GFP::RAB-3 localization 48hr post-axotomy in *ric-7* (A), *ric-7; dhc-1* (B), *ric-7; nud-2* (C) and *ric-7; jnk-1* (D) animals. Asterisk indicates cell body and bracket indicates dendrite. Scale bars = 10 μm. (E) Quantification of the number of dendritic RAB-3 puncta in intact and axotomized animals 48hr post-axotomy. Mean + SEM. **p<0.01; ****p<0.0001; ns, not significant. Unpaired t test. (F-I) Accumulation of GFP::RAB-3 at the tip of DA9 axons in intact *r/c-7* (F), *r/c-7; dhc-1* (G), *r/c-7; nud-2* (H) and *r/c-7; jnk-1* (I) animals (2d old adults). Magnified GFP channels are shown at the top right corner. Scale bars = 10 μm. (J) Quantification of GFP::RAB-3 intensity at the tip of axons of animals in (F-I). Mean + SEM. *p<0.05; ***p<0.0001. Unpaired t test. (K) Traces of the tail-bending behavior of *r/c-7* and *r/c-7; jnk-1* animals with and without axotomy 48hr post-axotomy. The shaded area indicates the 5 second stimulation. Mean + SEM. (L) Averaged tail angle during stimulation of the animals in (K). Numbers represent the number of animals. Mean and SEM. ***p<0.001; ****p<0.0001. Unpaired t test.

### JNK-1 modulates retrograde transport of SVs in response to axotomy

Since ectopic SV localization to the dendrite is triggered by axon injury, we next sought to find upstream mechanisms that link axonal injury to changes in SV localization. The best-characterized injury-sensing pathway in *C. elegans* is the DLK-1 MAP kinase pathway (Ghosh-Roy et al., 2010; Hammarlund et al., 2009; Nix et al., 2011; Yan and Jin, 2012; Yan et al., 2009), but SV localization to the dendrite after injury is independent of *dlk-1* (Figure 5B, 5D). We hypothesized that the JNK3 homolog *jnk-1* might play a role for two reasons (Kawasaki et al., 1999). First, *jnk-1* has been shown to be involved in the axon injury response in *Caenorhabditis elegans* GABA neurons. Loss of *jnk-1* results in improved morphological regeneration, while *jnk-1* overexpression inhibits morphological regeneration (Nix et al., 2014). Second, although *jnk-1* is not normally required for transport or localization of synaptic proteins in DA9, loss of *jnk-1* largely suppresses the synaptic phenotypes of *arl-8* mutants, suggesting that in some contexts *jnk-1* can regulate SV localization (Wu et al., 2013). We therefore examined whether *jnk-1* is involved in ectopic dendritic synapse formation in DA9 after axotomy. In uninjured neurons, SV distribution was largely unaffected by loss of *jnk-1*, with SV puncta confined to a specific region of the axon, and essentially absent from the dendrite (Figure 7 supplement 1C, 1E), consistent with previous results (Wu et al., 2013). By contrast, after axon injury *jnk-1* mutant animals showed very few SV puncta in the dendrite 48 hours (Figure 7D, 7E) and 12 hours after axotomy (Figure 7 supplement 1F, 1G). Thus, ectopic SV localization to the dendrite after axon injury requires the JNK3 homolog *jnk-1*.

Next, we investigated whether loss *of jnk-1* might alter SV localization by affecting dynein-dependent trafficking, as suggested by the similar phenotypes of the separate mutants (Figure 7B-7E). We analyzed the effect on SV localization in uninjured neurons, focusing on the distal tip of the DA9 axon. In control animals, SV accumulation was rarely seen at the axon tip (Figure 7F), indicating balanced anterograde and retrograde transport. However, in *nud-2*, *dhc-1*, and *jnk-1* mutant animals, especially the latter two, bright SV puncta were observed at the tip of almost every DA9 axon (Figure 7G-7J). These data suggest that even in uninjured neurons, *jnk-1* affects dynein-mediated traffic of synaptic vesicles. Thus, in injured neurons, *jnk-1* may function to promote dynein-driven SV movement into the dendrite, resulting in ectopic synapse formation.

### JNK-1 suppresses behavioral recovery after nerve injury

*jnk-1* mutant animals do not form ectopic synapses in the dendrite after injury. Since these ectopic synapses inhibit behavioral recovery after nerve injury, we tested the idea that *jnk-1* mutant animals might have improved behavioral recovery. Without axon injury, *jnk-1* mutants showed robust dorsal bending in response to light stimulation, although the degree of dorsal bending was smaller compared to controls (Figure 7K, 7L, *jnk-1*; *ric-7* uncut vs *ric-7* uncut, mean=59.7° and 70.1°, p°0.0003, unpaired t test). By contrast, 48 hours after axotomy, the *jnk-1* mutants showed improved dorsal bending behavior compared to controls (Figure 7K, 7L), similar to the improved behavior seen after dendrite removal (Figure 5F, 5G). Thus, altering neuronal cellular mechanisms to correctly direct information transfer in regenerated circuits improves behavioral recovery after nerve injury.

## Discussion

### Analyzing functional regeneration in DA9

This study establishes a new system for the study of axon regeneration in the DA9 neuron of *C. elegans*. We find that severing DA9 close to the dorsal nerve cord allows regeneration across the former synaptic area to occur in essentially all animals studied. Further, by using the *ric-7* genetic background to promote removal of the distal axon fragment, regenerative growth in DA9 occurs without confounding issues of axonal fusion or other potential effects of the remaining fragment. Previous studies in *C. elegans* have shown that the regenerating mechanosensory PLM neuron can reconnect to its distal axon segment after axotomy. This fusion event essentially restores the neuron back to its pre-injury state, without the need to rebuild synapses, and restores the full function of the circuit as measured by a light touch assay (Abay et al., 2017; Neumann et al., 2011). In this work, by using the *ric-7* background to promote distal segment removal, functional recovery can occur only if new, functional synapses are rebuilt onto relevant targets. In combination, our approach in DA9 in the *ric-7* background allows for the first time detailed morphological analysis of synapse reformation in a single neuron during axon regeneration. Further, by applying the modern neuroscience tools of calcium imaging and optogenetics to DA9 and its target muscles, it is possible to analyze the function of this single neuron in its endogenous circuit and also with respect to its ability to drive behavior. We expect these approaches to be useful for studies relating synapse formation to circuit function and behavior, both in the context of axon regeneration and also in uninjured neurons.

A potential concern is that loss of *ric-7* may cause phenotypes that interact with the question under study. *ric-7* encodes a protein of approximately 700 residues, with no clear vertebrate orthologs or functional domains. *ric-7* was first identified in a screen for genes involved in neuropeptide secretion. In *ric-7* mutants, neuropeptide secretion is reduced while acetylcholine release and response are unaltered (Hao et al., 2012). *ric-7* mutants were also found to have greatly enhanced degeneration of axon segments after axotomy, and this phenotype was found to be due to reduced localization of mitochondria to axons (Rawson et al., 2014). Interestingly, although mitochondria function is important for regeneration in *C. elegans* GABA neurons and in mammalian models (Cartoni et al., 2016; Han et al., 2016; Zhou et al., 2016), we found that *ric-7* animals have improved axon regeneration (Figure 1 supplement 1E-1H). Thus, in DA9 the loss of *ric-7* increases degeneration without negatively impacting regeneration. Further, we find that synaptic vesicle localization to the dendrite in response to injury is not dependent on *ric-7*, as it occurs at the same level in animals that are wild type and mutant at this locus (Figure 2H). Although the use of *ric-7* demands caution, our data indicate that it is a useful and effective tool for studying axon regeneration in DA9.

### Aberrant information rerouting limits behavior recovery after regeneration

We show that DA9 axons do regenerate and form functional synapses in the correct location after axotomy. However, we find that synapses also form ectopically in the dendrite, resulting in aberrant information routing and limiting behavioral recovery. Although ectopic synapse formation is reminiscent of the transformation of a dendrite to an axon after axon removal (Cho and So, 1992; Dotti and Banker, 1987; Fenrich et al., 2007; Gomis-Ruth et al., 2008; Hall et al., 1989; Hoang et al., 2005; Stone et al., 2010), we find that it is a fundamentally different process. First, dendrite-axon transformation requires axon removal, while in our experiments the axon is severed distant from the cell body, regenerates, and reforms functional synapses. Second, dendrite-axon transformation is accompanied by a rearrangement of MT polarity from dendritic orientation to axonal orientation (Stone et al., 2010; Takahashi et al., 2007). In the case of DA9, however, dendritic microtubule polarity as well as nAChR localization is maintained after axotomy (Figure 6F, Figure 6 supplement 1). Thus, our results define a novel cell-biological response to nerve injury.

The ability of dendritic synapses in DA9 to transfer information and cause specific behavioral defects likely depends on the circuit context within which DA9 is embedded. The dendrite of DA9 is in the ventral nerve cord, where it receives inputs from the AVA and AVD command neurons (Hall and Russell, 1991; White et al., 1986). What post-synaptic cells might respond to acetylcholine release from the DA9 dendrite to trigger ventral bending? The VNC contains a variety of ACh receptor fields. In particular, ventral muscles make synapses with ventral motor neurons in the VNC, such as VA and VB cells (White et al., 1986). Thus, the DA9 dendrite and ventral muscle receptor fields are both localized in the VNC. Our calcium imaging data indicate that DA9 activation can trigger ventral muscles (Figure 4B,4D), consistent with the idea that these muscles might respond directly to ACh release from DA9. However, it is also possible that information transfer from DA9 to ventral muscles is not monosynaptic. VNC motor neurons like VA and VB also have cholinergic receptor fields in the VNC, and could transfer excitatory signals from the DA9 dendrite to muscle. Thus, the particular anatomy of the *C. elegans* VNC helps determine the effect of dendritic release from DA9.

Synapses are highly organized junctions between pre-and post-synaptic cells, with vesicle release sites in tight apposition to post-synaptic receptors. In both DA9 axon and dendrite, postsynaptic cells may respond to synapse formation after DA9 injury. A key question for future study is to identify the direct synaptic targets of DA9 activity in axon and dendrite, and examine post-synaptic organization in these cells after injury. We found that the location of synapses after regeneration of the DA9 axon was largely similar to their initial location (Figure 2F). This result is reminiscent of key findings at the frog neuromuscular junction, in which denervation and reinnervation of muscle resulted in synapse formation at the original locations (Letinsky et al., 1976; Marshall et al., 1977; Rotshenker and McMahan, 1976). Thus, at least for DA9 axonal synapse regeneration, an attractive hypothesis is that agrin or other components of the extracellular matrix direct the location of synapses during regeneration, ensuring that they are correctly apposed to their post-synaptic partners.

### The role of JNK-1 in ectopic synapse formation

JNK, or c-Jun N-terminal kinase, is a MAP kinase that has been implicated in a variety of cellular processes such as immunity, stress response, tumor development and apoptosis (Davis, 2000). In both *Drosophila* and mouse, JNK signaling is required for efficient axon regeneration (Ayaz et al., 2008; Raivich et al., 2004). *C. elegans* has multiple JNK orthologs, including *kgb-1* and *jnk-1.* As in *Drosophila* and mouse, *kgb-1* is required for axon regeneration (Nix et al., 2011). By contrast, *jnk-1* suppresses axon regeneration, suggesting that different JNK pathways play different roles during regeneration (Nix et al., 2014).

Here we found that JNK-1 is required for ectopic synapse formation in the dendrite after axotomy (Figure 7D). Mutants in the dynein complex also rescued ectopic synapse formation (Figure 7B). In uninjured neurons, *jnk-1* mutants and dynein mutants both showed synaptic vesicle accumulation at the axon tip (Figure 7G-7J). These results suggest that JNK-1 functions to promote dynein-mediated SV transport. Thus, we propose that injury-activated JNK-1 affects the balance between anterograde and retrograde trafficking by promoting dynein-mediated SV transport. Consistent with this idea, loss of JNK-1, its upstream component JKK-1, or its scaffolding protein UNC-16/JIP3 causes SV mis-localization to the dorsal nerve cord in DD motor neurons at L1 stage (Byrd et al., 2001), and a similar phenotype is caused by mutations in the dynein complex (Kurup et al., 2017). In DA9, *jnk-1* has been shown to rescue the premature clustering of SVs in the asynaptic region caused by the *arl-8* mutation (Wu et al., 2013). This is also consistent with reduced dynein-mediated trafficking in the absence of JNK-1, which facilitates accumulation of SVs into the more distally localized synaptic region.

JNK-1 could regulate the dynein mediated transport of SVs in multiple ways. For example, activated JNK-1 could potentially phosphorylate cytoskeletal-interacting proteins, which could in turn influence motor processivity and vesicle transport (Brownlees et al., 2000; Chang et al., 2003; Otto et al., 2000; Reynolds et al., 2000). In addition, JNK interacting proteins (JIP) could also regulate cargo-motor or motor-motor interactions. Specifically, UNC-16/JIP-3 has been shown to act as a JNK-1 scaffolding protein and interact with both kinesin-1 and components of the dynein complex such as p150^glued^ and dynein light intermediate chain (Arimoto et al., 2011; Bowman et al., 2000; Cavalli et al., 2005). UNC-16/JIP-3 has also been implicated as an adaptor for dynein-mediated retrograde transport of activated JNK, which is important for transmission of damaging signals (Cavalli et al., 2005). It is possible that UNC-16 may regulate the activity of motors by bringing JNK-1 in close proximity to them after axonal injury. Some of these effects could also account for *jnk-1’s* effect on axon regeneration. Thus, loss of *jnk-1* could prevent ectopic synapse formation and also increase axon regeneration by affecting dynein-mediated transport.

### New synapses in the regenerating axon are not as efficient as intact ones

In our single-neuron system, although the regenerated synapses in the axon are able to drive some behavioral recovery, the level of dorsal tail bending is significantly lower than in intact controls. Even after removing the dendrite, which inhibits dorsal bending, full behavioral recovery is still not achieved. Since the number of regenerated synapses is similar to intact controls, individual regenerated synapses are likely less efficient than synapses in the intact neuron. Multiple effects could underlie this deficit, including defects in DA9 itself, defects in the post-synaptic muscle, imprecise alignment of the two cells, or a combination of these factors. Detailed morphological and molecular characterization of regenerated synapses is needed to understand the difference between them and intact synapses. Our study highlights the difficulty in restoring normal circuit function after nerve injury, and provides insight into specific cellular choke points that need to be resolved to achieve complete recovery.

## Materials and Methods

### Strains

Animals were maintained at 20°C on NGM plates seeded with 0P50 *E. coli* according to standard methods. The following strains were obtained from the *Caenorhabditis* Genetic Center (CGC): MT6924 [*ric-7(n2657)*], NM1489 [*dhc-1(js319); jsIs37*], VC8 [*jnk-1(gk7)*], RB1022 [*nud-2(ok949)*], CZ5730 [*dlk-1(ju476)*]. TV12772 [*wyIs386(Pitr-1::acr-2::gfp)*] was provided by the Yogev lab.

The following transgenes were generated by microinjection (Promega 1kb DNA ladder was used as a filler in the injection mixes): *wpIs101 [Pitr-1 pB::gfp::rab-3::SL2::mcherry@50ng/uL; Pmyo-2::mcherry@2ng/uL*], *wpIs102* [*Pitr-1 pB::gfp::rab-3::SL2::mcherry@50ng/uL; Pmyo-2::mcherry@2ng/uL*], *wpIs98* [*Pitr-1 pB::gfp::Chrimson::SL2::mcherry@20ng/uL; Podr-1::rfp@30ng/uL*]. *wpIs103* [*Pmyo-3::GCaMP6@80ng/uL; Pmyo-2::mcherry@2ng/uL*], *wpEx294* [*Pitr-1 pB::ebp-2::gfp::SL2::mcherry@50ng/uL; Pmyo-2::mcherry@2ng/uL*] and *wpEx295* [*Pitr-1 pB::mcherry@25ng/uL; Pmyo-2::mcherry@2g/uL*].

### List of strains used in this study

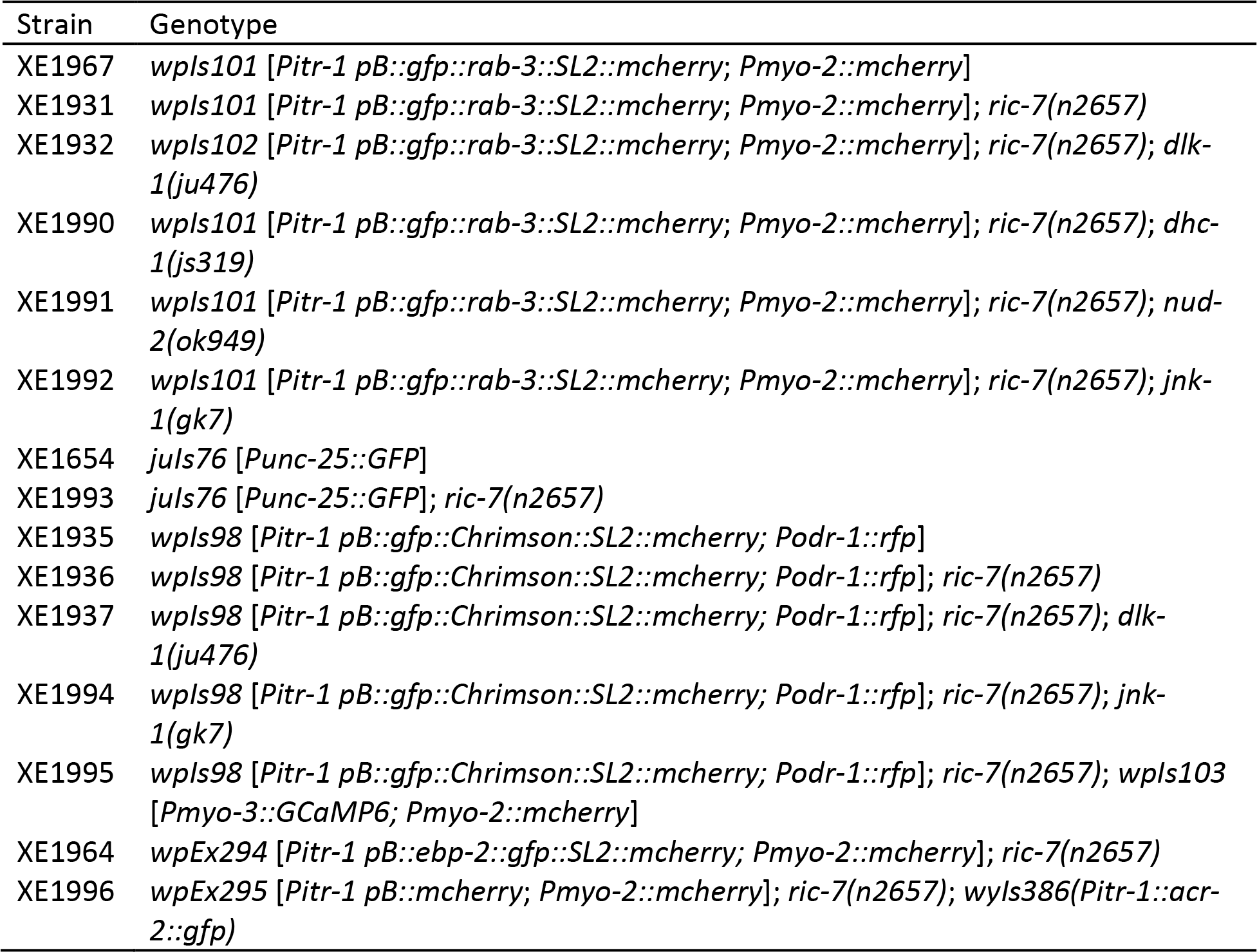

| Strain | Genotype |
| --- | --- |
| XE1967 | *wpls101* [*Pitr-1 pB::gfp::rab-3::SL2::mcherry*; *Pmyo-2::mcherry*] |
| XE1931 | *wpls101* [*Pitr-1 pB::gfp::rab-3::SL2::mcherry*; *Pmyo-2::mcherry*]; *ric-7(n2657)* |
| XE1932 | *wpls102* [*Pitr-1 pB::gfp::rab-3::SL2::mcherry*; *Pmyo-2::mcherry*]; *ric-7(n2657)*; *dlk-1(ju476)* |
| XE1990 | *wpls101* [*Pitr-1 pB::gfp::rab-3::SL2::mcherry*; *Pmyo-2::mcherry*]; *ric-7(n2657)*; *dhc-1(js319)* |
| XE1991 | *wpls101* [*Pitr-1 pB::gfp::rab-3::SL2::mcherry*; *Pmyo-2::mcherry*]; *ric-7(n2657)*; *nud-2(ok949)* |
| XE1992 | *wpls101* [*Pitr-1 pB::gfp::rab-3::SL2::mcherry*; *Pmyo-2::mcherry*]; *ric-7(n2657)*; *jnk-1(gk7)* |
| XE1654 | *juls76* [*Punc-25::GFP*] |
| XE1993 | *juls76* [*Punc-25::GFP*]; *ric-7(n2657)* |
| XE1935 | *wpls98* [*Pitr-1 pB::gfp::Chrimson::SL2::mcherry*; *Podr-1::rfp*] |
| XE1936 | *wpls98* [*Pitr-1 pB::gfp::Chrimson::SL2::mcherry*; *Podr-1::rfp*]; *ric-7(n2657)* |
| XE1937 | *wpls98* [*Pitr-1 pB::gfp::Chrimson::SL2::mcherry*; *Podr-1::rfp*]; *ric-7(n2657)*; *dlk-1(ju476)* |
| XE1994 | *wpls98* [*Pitr-1 pB::gfp::Chrimson::SL2::mcherry*; *Podr-1::rfp*]; *ric-7(n2657)*; *jnk-1(gk7)* |
| XE1995 | *wpls98* [*Pitr-1 pB::gfp::Chrimson::SL2::mcherry*; *Podr-1::rfp*]; *ric-7(n2657)*; *wpls103* [*Pmyo-3::GCaMP6*; *Pmyo-2::mcherry*] |
| XE1964 | *wpEx294* [*Pitr-1 pB::ebp-2::gfp::SL2::mcherry*; *Pmyo-2::mcherry*]; *ric-7(n2657)* |
| XE1996 | *wpEx295* [*Pitr-1 pB::mcherry*; *Pmyo-2::mcherry*]; *ric-7(n2657)*; *wyls386(Pitr-1::acr-2::gfp)* |

### Cloning and Constructs

Plasmids were assembled using Gateway recombination (Invitrogen). Entry clones were generated by Gibson Assembly when needed.

### Laser Axotomy

Laser axotomy was performed as described previously (Byrne et al., 2011). In brief, L4 stage animals were immobilized with 0.1 μm diameter polystyrene beads (Polysciences) and mounted on a 3% agarose pad on a glass slide. The animals were visualized with a Nikon Eclipse 80i microscope using a 100× Plan Apo VC lens (1.4 NA). DA9 axons were cut at the posterior part of the asynaptic region using a 435nm Micropoint laser by 10 pulses at 20Hz. In some experiments DA9 dendrites were cut near the cell body or the DA9 cell body was abated. For the axotomy of GABA motor neurons, the first, fourth and seventh axons from the tail were cut at the midpoint of the commissures. Animals were then recovered to 0P50 seeded NGM plates and analyzed later.

### Fluorescence Microscopy and Quantification

To assess DA9 axonal and synaptic regeneration, animals were imaged at different time points after axotomy. Animals were immobilized with 4-40 mM levamisole and mounted on a 3% agarose pad on a slide. Images were then acquired as 0.3 μm z stacks on a spinning-disc confocal microscope (PerkinElmer UltraVIEW VoX, Nikon Ti-E Eclipse inverted scope, Hamamatsu C9100-50 camera) with a CFI Plan Fluor 40X oil objective (1.3 NA) using Volocity software (PerkinElmer). Throughout this study, all images were acquired with the same exposure time, camera sensitivity and laser power. Images were then exported as tiff files and analyzed in ImageJ.

Maximum intensity projection was created from the tiff files for analysis. DA9 axon or dendrite length was measured using an ImageJ plugin, Simple Neurite Tracer (v3.1.0). Injury-induced degeneration was quantified manually with beadings, breaks and clearance of the distal axon fragment as the criteria. To count synaptic vesicle puncta in the DA9 axon, an ROI around the axon was drawn. A default threshold was determined by ImageJ based on the intensity histogram and applied to the ROI. This created a binary mask of SV puncta in the axon. The mask was then processed with the Watershed function in order to separate puncta close to each other. Finally, the number of puncta was measured using the Analyze Particle function in ImageJ. The results were further confirmed manually. This thresholding formula provided a compromise between including all puncta and avoiding background noise.

A different method was used to count SV puncta in the dendrite because in uncut animals the SV signal was extremely weak, so automatic thresholding created artifacts. Briefly, line scanning was performed along the dendrite, and a uniform threshold of 2500 fluorescent intensity was applied. Peaks above the threshold were counted as SV puncta.

To measure axon regeneration of GABA motor neurons, animals were immobilized with polystyrene beads on a 3% agarose pad and imaged at 24 hours after axotomy. Images were acquired using a UPLFLN 40X oil objective (1.3 NA) on an Olympus BX61 microscope equipped with an Olympus DSU and a Hamamatsu ORCA-Flash4.0 LT camera. Acquisition was controlled using the MicroManager software.

Images were analyzed in ImageJ. Briefly, GABA axon length along the dorsal-ventral body axis was measured and normalized to the full axis length. The distribution of the normalized length was used to compare regeneration.

### Optogenetics and Behavior

Animals expressing Chrimson in DA9 were cultured with 0P50 containing 10 mM all-trans-retinal (ATR) on NGM plates. Prior to the behavior assay, animals were transferred to a fresh plate without 0P50. The plate was then placed under a Leica M165FC stereo scope. Movies were taken with a Basler acA2440 camera controlled by the WormLab software. A Prior Lumen 200 fluorescent source and a Leica mCherry filter set were used to give light stimulation. Light intensity was about 300 W/m^2^ throughout the study. Bright field illumination was kept at a low intensity to reduce non-specific activation of Chrimson. The tail bending behavior was analyzed using the Wormlab software. In brief, the animal was detected by thresholding, and then the body midline was segmented evenly with 14 points. The angle between the last three points was used as the tail angle.

### Calcium Imaging

Animals expressing Chrimson in DA9 and GCaMP6 in BWMs were cultured with 0P50 and ATR. Animals were immobilized by polystyrene beads and mounted on 3% agarose pads on slides. Movies were taken on an Olympus BX61 microscope with a Hamamatsu 0RCA-Flash4.0 LT camera. An X-Cite XLED1 was used as the light source (EXCETILAS). Specifically, a blue LED module (BDX (450-495nm)) was used to image GCaMP6 and a red LED module (RLX(615-655nm)) was used to give continuous stimulation.

To reduce the activation of Chrimson by blue light, a ND4.0 filter was mounted in front of the BDX module and the power of blue light was set to 10%. Images were acquired at 5Hz using the HCImageLive software. Movies that had a stable baseline before stimulation were used for analysis in ImageJ. Briefly, an ROI was selected far away from the animal to generate a background fluorescence intensity, Fb. ROIs of the posterior dorsal and ventral BWMs were also drawn to detect Ca^2+^ signals, F_t_. F’_t_=Ft-Fb was used to calculate Ca^2+^ changes. Then the baseline, F0, was calculated as averaged F_t_’ from the first 5s of the movie. Finally, F=(F’_t_-F_0_)/F_0_ was used to display Ca^2+^ signal changes.

### Time-lapse Imaging and quantification

Time-lapse imaging was performed to visualize EBP-2::GFP traces and SV trafficking in DA9. Animals were immobilized by 4-40mM levamisole and mounted on 3% agarose pads on slides. Movies were taken on an Olympus BX61 microscope with a Hamamatsu ORCA-Flash4.0 LT camera at 5 or 10 Hz for 1 min. Movies were then analyzed in ImageJ. Briefly, a line was drawn along the dendrite or the axon and the kymograph was generated using the Reslice function. EBP-2 and SV traces were analyzed from the kymograph and confirmed manually in the movie.

## Acknowledgements

We thank the international *C. elegans* Gene Knockout Consortium, the *Caenorhabditis* Genetic Center and the Yogev lab for strains. We thank the Colón-Ramos lab for sharing the Chrimson and GCaMP6 plasmids. We thank everyone in the Hammarlund lab for feedback and suggestions.

---

**Figure 1 supplement 1.**
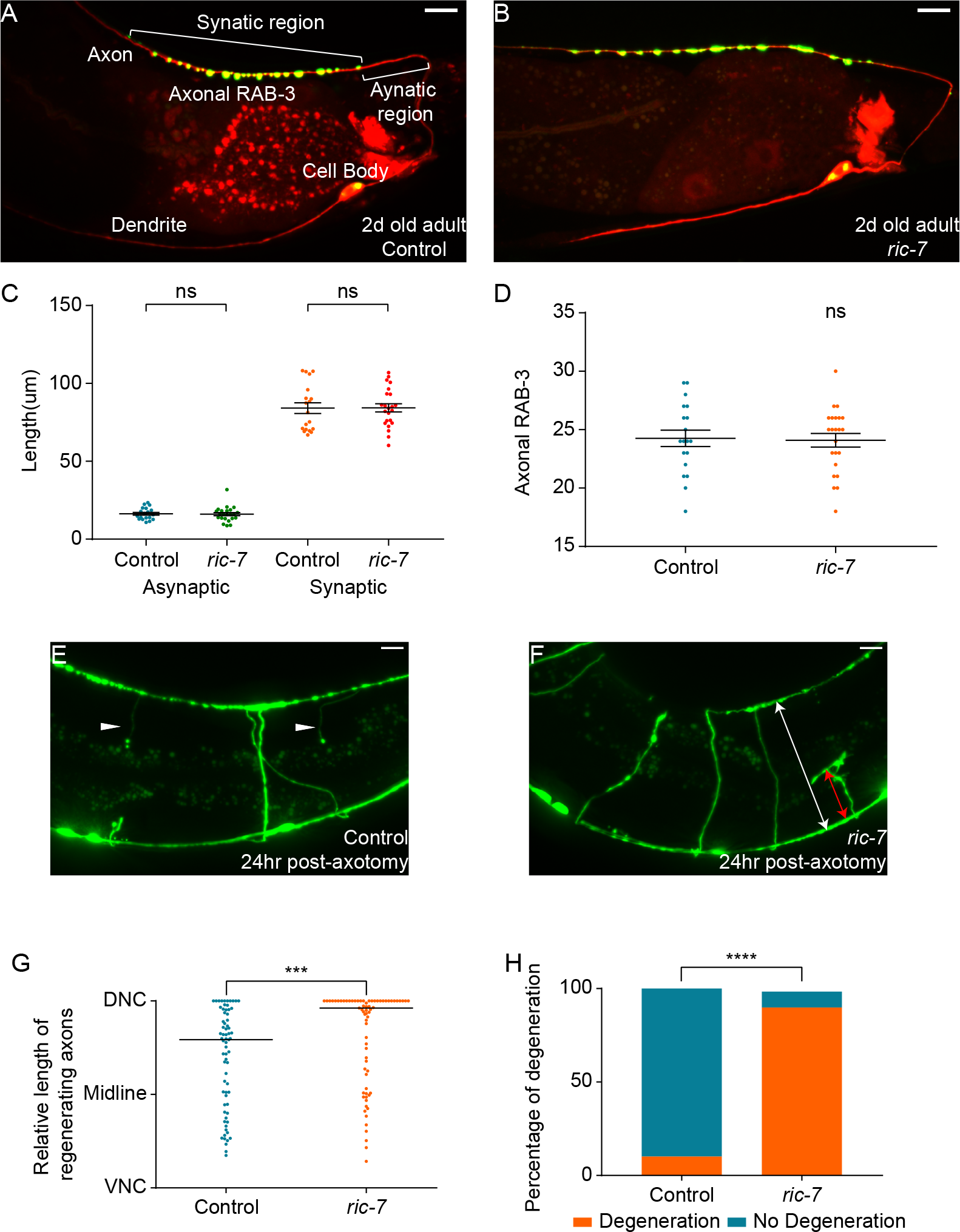
*ric-7* Animals Show Normal DA9 Development, Better Regeneration and Increased Degeneration of the GABA Motor Neurons. (A and B) GFP::RAB-3 and mCherry labeling in DA9 of intact control (A) and *ric-7* (B) animals (2d old adults). Scale bars = 10 μm. (C) Length of the asynaptic region and the synaptic region in intact control and *ric-7* animals (2d old adults). Mean ± SEM. ns, not significant. Unpaired t test. (D) Number of axonal RAB-3 puncta in intact control and *ric-7* animals (2d old adults). Mean + SEM. ns, not significant. Unpaired t test. (E and F) Regeneration and degeneration of GABA motor neurons in control (E) and *ric-7* (F) animals 24hr post-axotomy. White arrowheads point to the remaining distal axon segments in control (E) which have degenerated in *ric-7* (F). The red line in (F) indicates the length of the regenerating axon along the dorsal-ventral axis and the white line indicates the width of the animal. Scale bars = 10 μm. (G) Relative length of regenerating axons (the ratio of the length of red line and white line in (F)) in control and *ric-7* animals 24hr post-axotomy. Black line is median. ***p<0.001. Kolmogorov-Smirnov test. (H) Percentage of degenerating axons of GABA motor neurons in control and *ric-7* animals 24hr post-axotomy. ****p<0.0001. X^2^ test.

**Figure 3 supplement 1.**
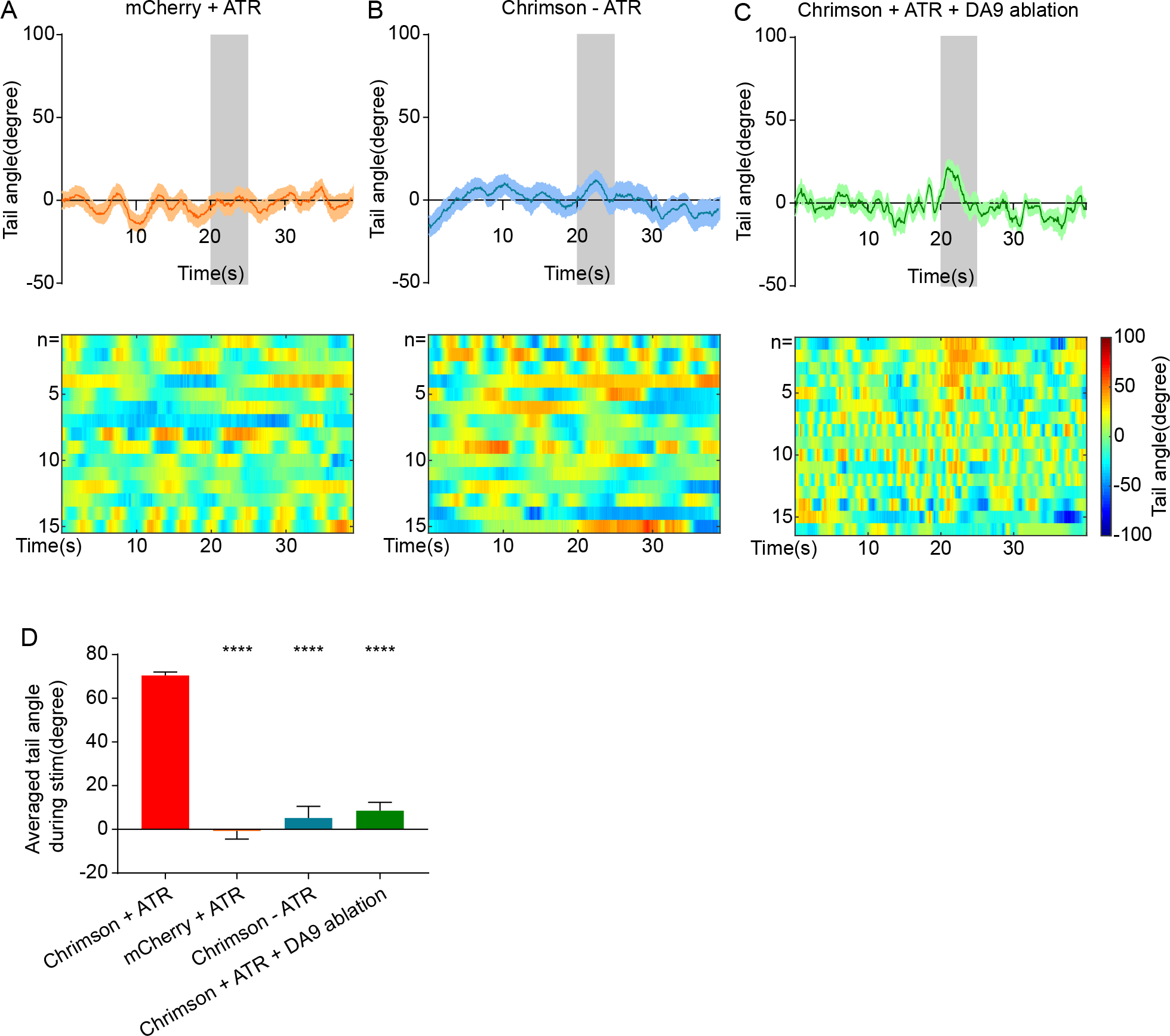
Optogenetically Triggered Tail-bending Behavior Requires Chrimson and ATR and is DA9 Specific. (A) *ric-7* animals expressing mCherry instead of Chrimson supplied with ATR show no dorsal tail bending. Mean + SEM. The shaded area represents the 5 seconds stimulation. (B) *ric-7* animals expressing Chrimson in the absence of ATR show no dorsal tail bending. Mean + SEM. The shaded area represents the 5 seconds stimulation. (C) WT animals expressing Chrimson supplied with ATR but with their DA9 cell body ablated show no dorsal tail bending. Mean + SEM. The shaded area represents the 5 seconds stimulation. (D) Averaged tail angle during stimulation of positive controls and animals in (A), (B) and (C). The positive controls are the same as in Figure 3C. Mean and SEM. ****p<0.0001. Unpaired t test.

**Figure 5 supplement 1.**
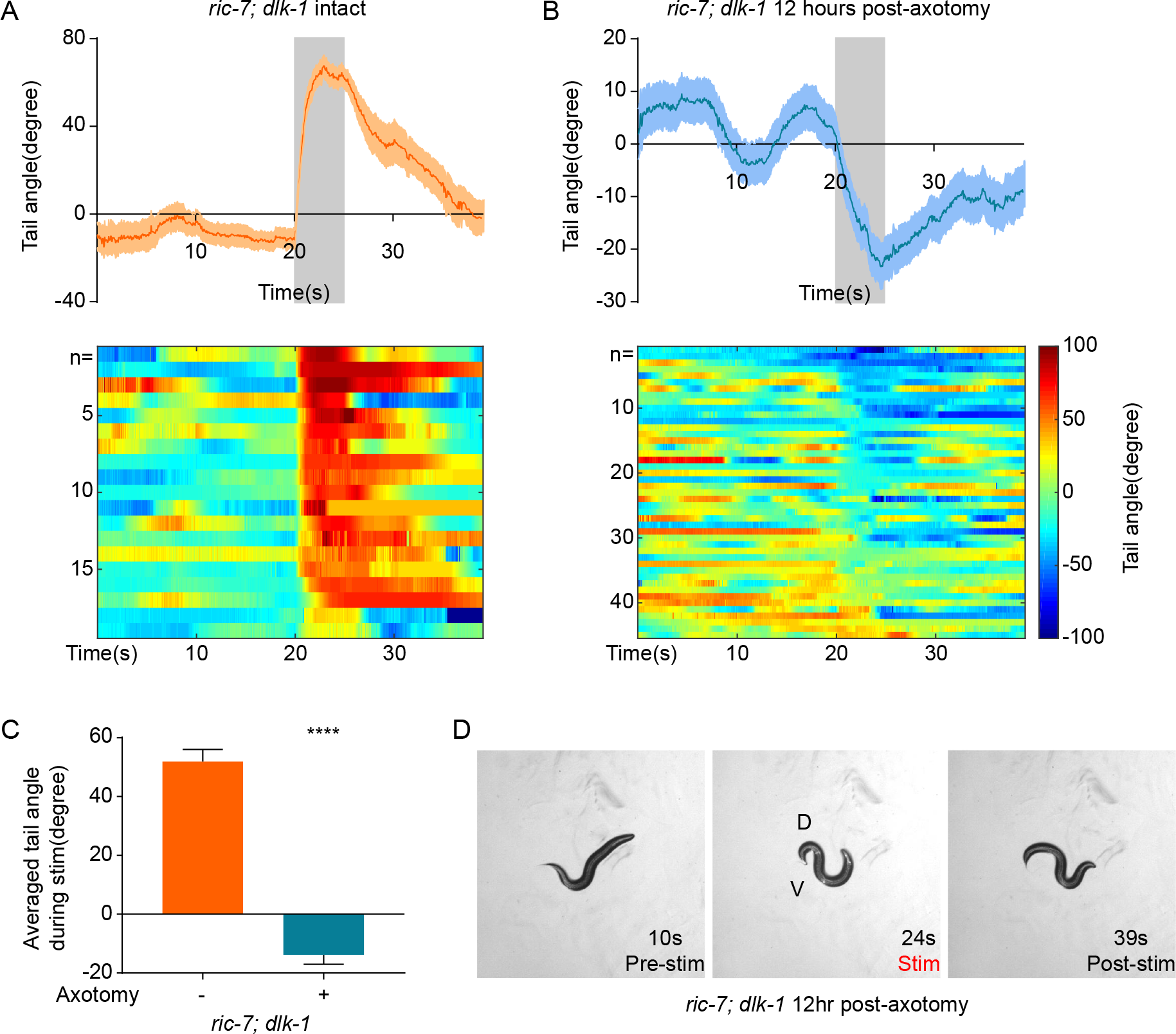
Behavior Response of *ric-7*; *dlk-1* Animals with and without Axotomy. (A) *ric-7*; *dlk-1* intact animals show robust dorsal tail bending. The shaded area represents the 5 seconds stimulation. Mean + SEM. (B) *ric-7; dlk-1* axotomized animals show robust ventral tail bending 12hr post-axotomy. Mean + SEM. (C) Averaged tail angle during stimulation of animals in (A) and (B). Mean and SEM. ****p<0.0001. Unpaired t test. (D) Images from a movie showing the ventral-bending behavior of a *ric-7; dlk-1* animal 12hr post-axotomy.

**Figure 6 supplement 1.**
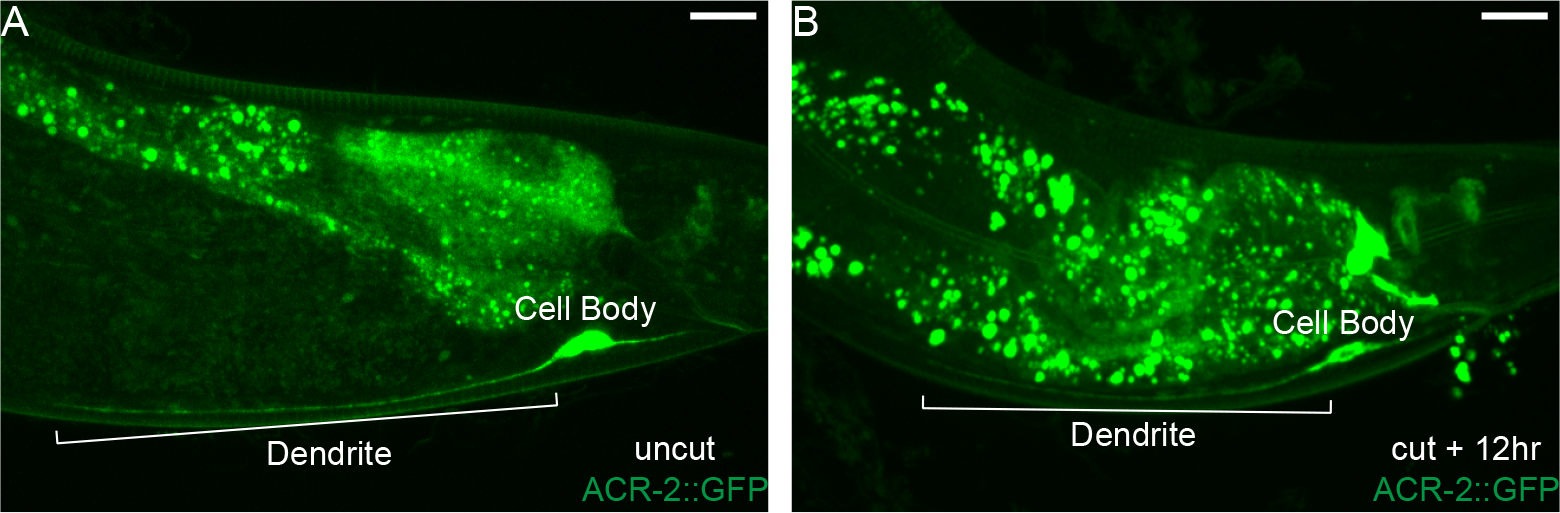
ACR-2, a Subunit of the Nicotinic Acetylcholine Receptor, Maintains Dendritic Localization post-Axotomy. (A) ACR-2::GFP localizes to the dendrite and soma of DA9 in intact *ric-7* animals. The scale bar = 10 μm. ACR-2::GFP still localizes to the dendrite and soma of DA9 in axotomized *ric-7* animals 12hr post-axotomy. The scale bar = 10 μm.

**Figure 7 supplement 1.**
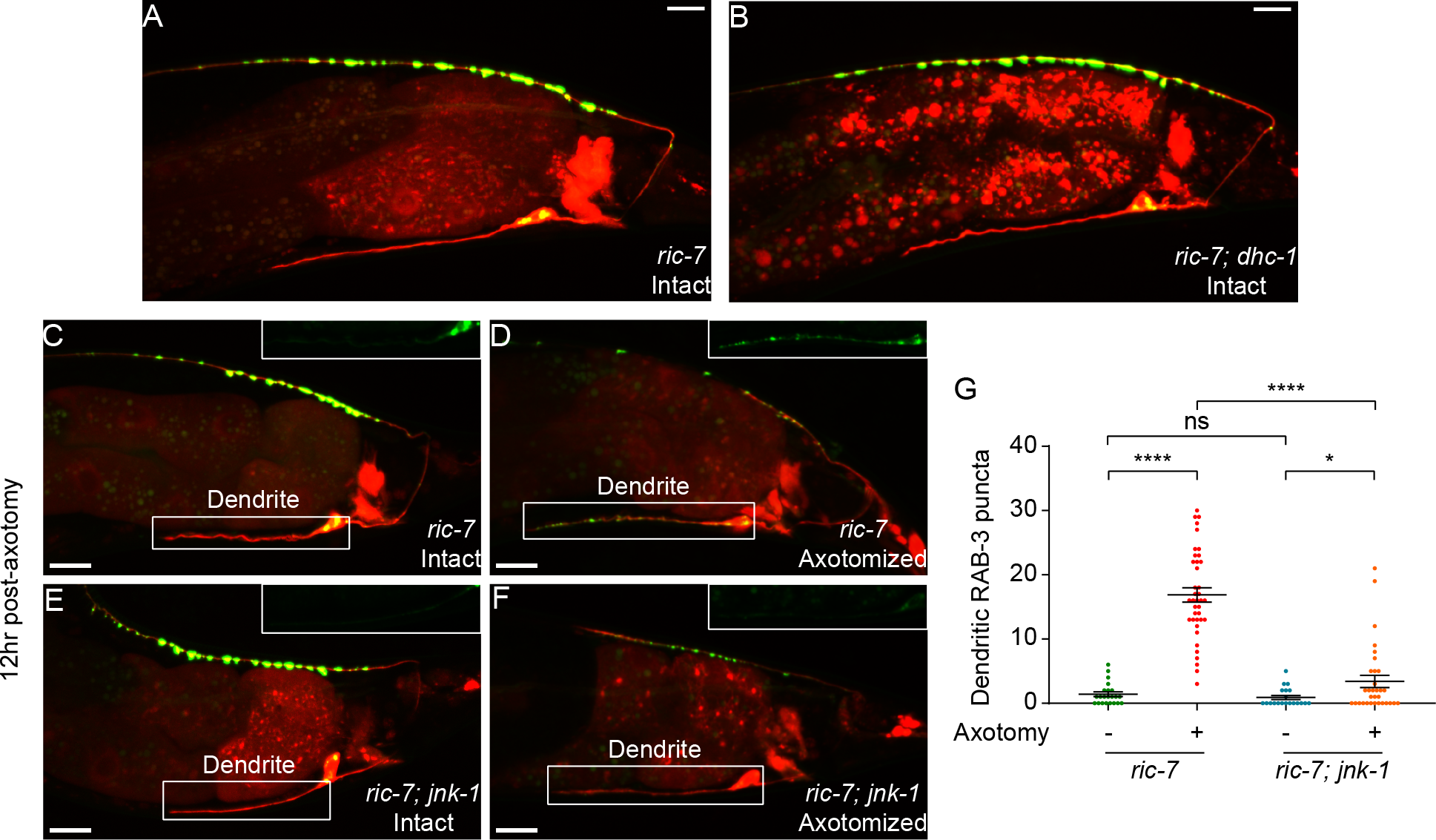
DA9 in *dhc-1* Dnimals Develops Normally and Loss of JNK-1 Rescues RAB-3 Mis-localization in DA9 Dendrite 12 hr post-Axotomy. (A-B) RAB-3::GFP and mCherry labeling of DA9 in intact *ric-7* and *ric-7; dhc-1* animals (2d old adults). Scale bars = 10 μm. (C-F) RAB-3 localization to the dendrite in *ric-7* and *ric-7; jnk-1* animals with and without axotomy 12hr post-axotomy. Magnified GFP channels are shown at the top right corner. Scale bars = 10 μm. (G) Quantification of dendritic RAB-3 puncta number in animals in (C-F). Mean + SEM. *p<0.05; ****p<0.0001; ns, not significant. Unpaired t test.

